# Long-term monitoring of inflammation in the mammalian gut using programmable commensal bacteria

**DOI:** 10.1101/075051

**Authors:** David T Riglar, Michael Baym, S Jordan Kerns, Matthew J Niederhuber, Roderick T Bronson, Jonathan W Kotula, Georg K Gerber, Jeffrey C Way, Pamela A Silver

## Abstract

Inflammation in the gut, caused by infection and autoimmunity, remains challenging to effectively detect, monitor, and treat. Here, we engineer a commensal mouse *E. coli* strain to record exposure to tetrathionate, a downstream product of reactive oxygen species generated during inflammation. Using these programmed bacteria to sense *in situ* levels we show that tetrathionate accompanies inflammation during Salmonella-induced colitis in mice and is elevated in an inflammatory bowel disease mouse model. We demonstrate long-term genetic stability and associated robust function of synthetic genetic circuits in bacteria colonizing the mammalian gut. These results demonstrate the potential for engineered bacteria to stably and reliably probe pathophysiological processes for which traditional diagnostics may not be feasible or cost-effective.

**One sentence summary:** Engineered bacteria record an inflammatory response in an IBD mouse model and are genetically stable during long-term growth in the mouse gut.

## Main Text

The development of bacteria capable of robust diagnostic and therapeutic action in the complex intestinal environment has been a long-held goal of synthetic biology. To this end, progress has been made in engineering synthetic genetic circuits to retain sustained memory of transient gene expression in bacteria under laboratory ^1-4^ and short-term *in vivo* settings ^5, 6^. The long-term stability and function of such circuits has never been determined in more realistic *in vivo* settings.

Monitoring and near real-time treatment of inflammatory gut conditions, such as inflammatory bowel disease (IBD) is a high priority ^7^. For example, the emerging concept in IBD management of “treating to target” until no subclinical inflammation remains ^8^, requires routine monitoring of inflammation. Unfortunately, the currently available indirect inflammatory biomarkers used in the gut (serum C-reactive protein, fecal calprotectin and stool lactoferrin) have sensitivity and specificity trade offs that maintain clinical reliance on the endoscopic gold standard ^7^. The high cost, difficulty for quantitative interpretation ^9, 10^, risk of complications and low availability of endoscopy, however, make it inappropriate for routine monitoring.

Many intestinal inflammatory responses cannot be accurately assessed using traditional non-invasive diagnostics such as fecal testing. This is in part due to localized responses involving transient chemical species that either passively degrade or are actively utilized by the native microbiota, or the host, prior to excretion in feces. For example, activated neutrophils produce short-lived superoxide and downstream reactive oxygen and nitrogen species (ROS/RNS) ^11^.

The conversion of thiosulfate into tetrathionate downstream of ROS in the mouse intestine has been described during *Salmonella typhimurium* ^12^ and *Yersinia enterocolitica* ^13^ induced inflammation (Figure S1). Tetrathionate, sensed through the TtrR/TtrS two-component system ^14^, provides a growth advantage to these pathogens during inflammation ^13, 15, 16^ and may also be utilized by other microbes, particularly pathogens ^17^. Elevated tetrathionate levels in toll-like receptor 1 deficient mice ^13^ and the enrichment of tetrathionate utilization genes in the microbiota of a Tbet^−/−^ Rag^−/−^ mouse ulcerative colitis (TRUC) model ^18^ point to the molecule’s potential as a more general marker of inflammation, including during IBD. However, as detection is not possible by non-invasive methods this has prohibited further investigation, particularly during human disease.

Here we engineer a tetrathionate-responsive commensal *E. coli* strain, NGF-1, based on our previous system for antibiotic detection and memory ^5^. The system contains two parts: 1) the ‘trigger’, an environmentally-responsive promoter driving Cro protein expression (Figure 1A, upper); and 2) the ‘memory element’, a Cro-inducible CI/Cro transcriptional switch derived from phage lambda ^5^ (Figure 1A, lower). Once memory is initiated, expression of a *lacZ* reporter gene allows memory-state quantification using indicator culture plates (Figure 1A, lower). We utilized *ttrR/S* genes and the P_ttrBCA_ promoter from *S. typhimurium* to drive Cro expression (Figure 2A, upper) creating strain PAS638 (Figure 1A; Table S1).

**Figure 1:**
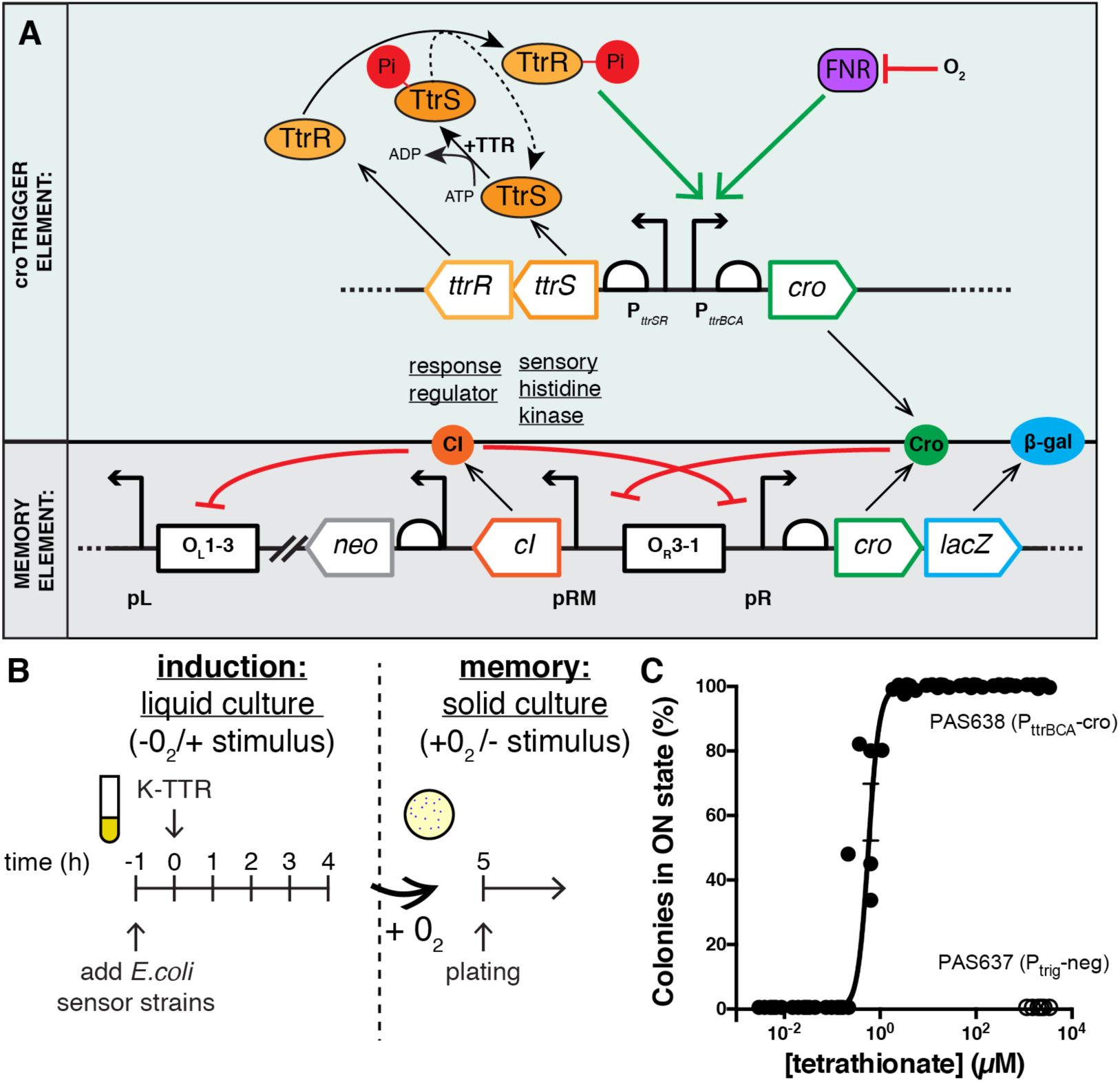
Engineering and testing a tetrathionate responsive memory device in *E.coli* NGF-1. A) A bacterial memory device, PAS638, was constructed in mouse commensal *E. coli* NGF-1. *S. typhimurium ttrR/S* and P_ttrBCA_ drive Cro ‘trigger’ expression to switch a phage lambda-based memory circuit ^5^. In the presence of tetrathionate, TtrS becomes phosphorylated, in turn phosphorylating TtrR, which activates expression through P_ttrBCA_ in anaerobic conditions. Cro protein expression switches memory ON, accompanied by *lacZ* reporter expression. B) Testing of *in vitro* memory showed. C) a dose-response curve of PAS638 (EC50: 0.38-0.85µM 95%CI) but no response of the triggerless control PAS637 strain. Graph shows individual values from 6 biological replicates, non-linear fit ± SEM.

**Table 1:**
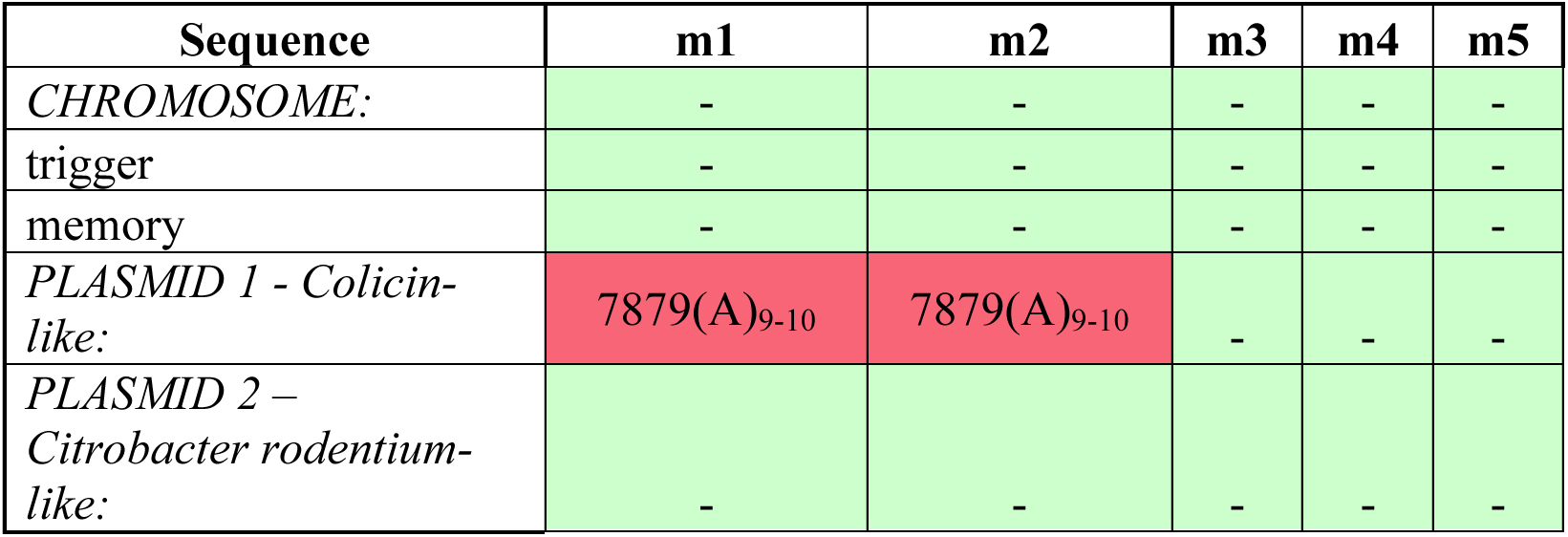
PAS638 does not accrue mutations in synthetic elements over long periods colonizing the mammalian gut. Mutational analysis of whole genome sequencing results from PAS638 following colonization of the mouse for 159 days showed no mutation in chromosomal or the *Citrobacter rodentium* – like plasmid. One mutational insertion in the Colicin-like plasmid was identified in two cohoused mice, m1 and m2.

**Figure 2:**
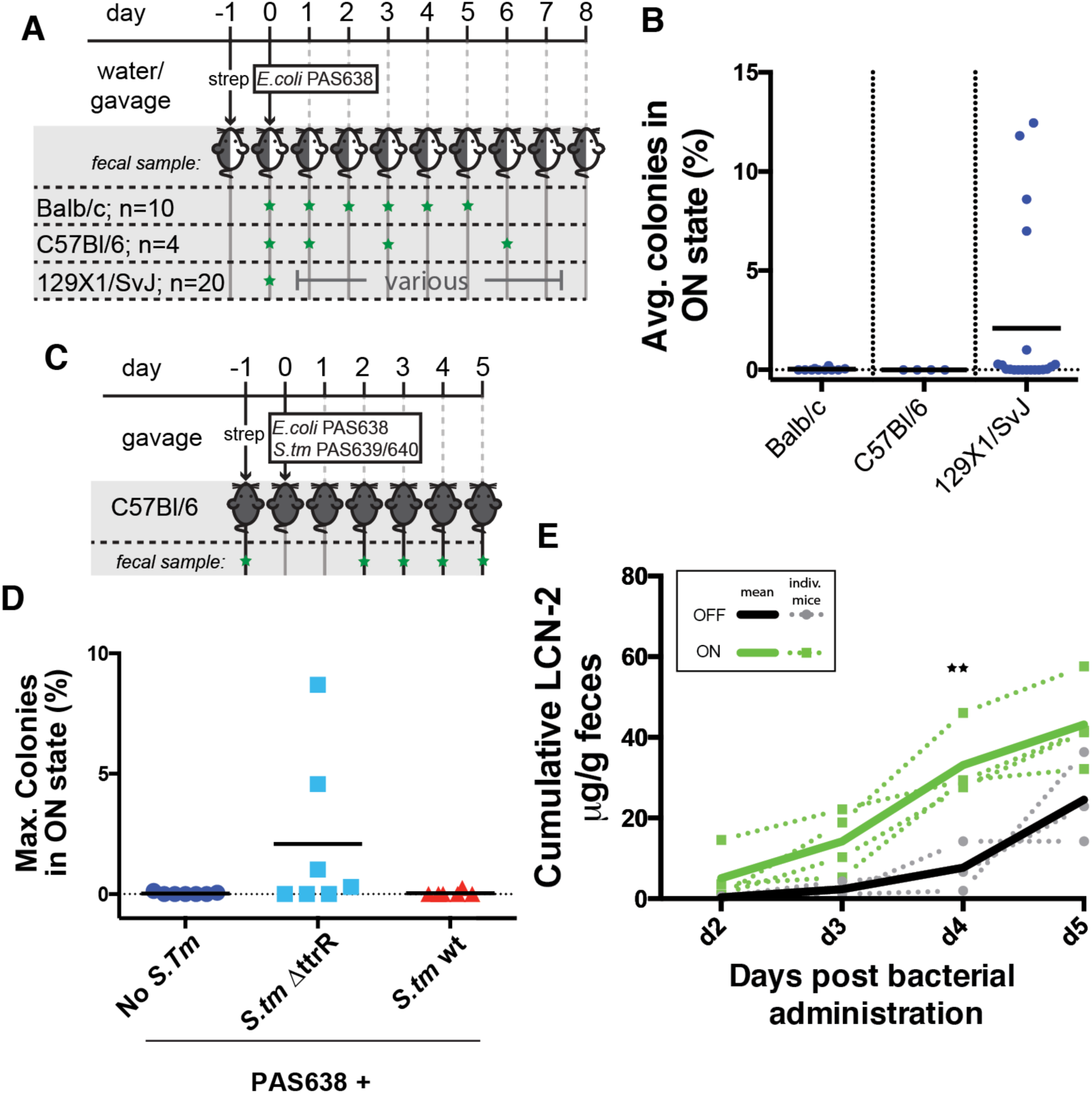
Tetrathionate-responsive memory bacteria specifically sense tetrathionate and correspond to severity of inflammation *in vivo*. A) Balb/c, C57Bl/6 and 129X1/SvJ mice were colonized with PAS638 and the bacterial memory state was analyzed (green star = fecal sample collected). B) Balb/c and C57Bl/6 mice showed no switching, while a subset of 129X1/SvJ mice showed elevated response. Graph shows averages from at >3 days for individual mice and means. C) PAS638 and *S. typhimurium* 14028s *(S. tm)* bacteria were administered by oral gavage to mice one day after streptomycin treatment, and fecal samples were analyzed d2-5 post administration (green stars) D) showing specific response of PAS638 on day 4 and/or 5 (see Figure S2A) when co-infected with *S. typhimurium* ΔttrR bacteria but not control or *S. typhimurium* wt bacteria. E) Cumulative LCN-2 levels in mice administered PAS638+ *S. typhimurium* ΔTtrR were higher in mice with PAS638 response (green lines) than in those without (black lines). Graph shows plots for individual mice (dotted) and ON or OFF averages (solid lines). * p<0.01 using multiple t-tests with Holm-Sidak multiple comparisons test and each timepoint analyzed individually without assuming a consistent SD.

This engineered strain exhibits memory upon tetrathionate exposure. We induced memory under anaerobic liquid culture conditions to trigger P_ttrBCA_ promoter expression in the presence of tetrathionate and quantified memory on aerobically grown plates in the absence of tetrathionate, which turned P_ttrBCA_ expression off (Figure 1B). PAS638 bacteria showed a characteristic dose-response curve upon tetrathionate induction (EC50:0.38-0.85µM 95% CI) (Figure 1C). By comparison, the triggerless control strain, PAS637 (Table S1), showed no switching at 2mM tetrathionate (Figure 1C). Previous *in vivo* quantification did not detect tetrathionate in inflammation-free wild-type mice but showed concentrations in a low millimolar range under inflammatory conditions, suggesting the sensitivity of PAS638 is well adapted to physiologically relevant concentrations of the analyte ^12, 13^.

PAS638 showed no tetrathionate response in two mouse strains and low-level response in a third, indicating the general absence of tetrathionate in healthy mice. For these baseline experiments we administered PAS638 to Balb/c, C57Bl/6 or 129X1/SvJ mice. We measured response through screening at least three serial fecal samples collected over the subsequent 5-8 days (Figure 2A). Results showed no notable response of PAS638 in Balb/c or C57Bl/6 mice, but bacteria from a subset of 129X1/SvJ mice showed elevated switching (Figure 2B). We conclude PAS638 is not sensitive to background levels of tetrathionate in healthy Balb/c or C57Bl/6 mice. The elevated response in 129X1/SvJ mice was surprising, however, may result from known defects in macrophage recruitment to sites of inflammation in this strain ^19, 20^. Based on this result, testing of the strains specificity for tetrathionate was undertaken in a C57Bl/6 background.

We tested the ability of PAS638 to detect inflammation during intestinal infection using murine *S. typhimurium* colitis as a model ^21^. *S. typhimurium* strains that are tetrathionate reduction deficient cause elevated cecal tetrathionate, while strains with wild-type reduction capabilities do not ^12^. To test the specificity of PAS638, we pre-treated C57Bl/6 mice with streptomycin by oral gavage, and administered them PAS638, PAS638 with *S. typhimurium,* or PAS638 with a *S. typhimurium* ΔttrR variant which is unable to express tetrathionate reductase (Figure 2C; Table S1) ^16^. We screened fecal samples collected on days 2-5 post-administration and observed no tetrathionate sensing in samples from control and *S. typhimurium* infected mice (Figure 2D). However, PAS638 co-administered with *S. typhimurium* ΔttrR showed elevated switching in a subset of mice (Figure 2D) with switching occurring on days 4-5 post-infection (Figure S2A). Titers of PAS638 *E. coli* (Figure S2B) and *S. typhimurium* variants (Figure S2C) were consistent between groups, demonstrating that variability in colonization was not responsible for the differences observed.

We also evaluated inflammation by lipocalin-2 (LCN-2) biomarker quantification (Figure S2D) ^22^ and by blinded histopathology scoring of cecum and colon sections (Figure S2E-F). Together, these techniques identified inflammation in all *S. typhimurium* and *S. typhimurium* ΔttrR infected mice ^12^. Of particular note, when mice from the PAS638 + *S. typhimurium* ΔttrR group were analyzed independently, mice in which PAS638 colonies in the on state had been detected had significantly higher (p<0.01) cumulative LCN-2 values at day 4 post-infection than those in which PAS638 remained off (Figure 2E). The increasing rate of LCN-2 level rise at day 5 of the experiment suggests that longer-term monitoring of PAS638 switching would be ideal, however, this was not possible in the model due to susceptibility of C57Bl/6 mice to systemic salmonellosis. Nevertheless, these results demonstrate that our engineered memory strain specifically senses the presence of tetrathionate and that tetrathionate sensing generally corresponds to a more acute inflammatory response *in vivo*.

Increased ROS levels are associated with IBD ^23^. Interleukin-10 deficient mice (IL10^−/−^) share characteristics with human IBD, including influence of the microbiota composition on disease progression ^24^. To test the ability of PAS638 to detect subclinical inflammation in an IBD context, we colonized ~6 month-old, retired-breeder IL10^−/−^ mice that were not expected to have active colitis (due to being raised in gnotobiotic and barrier SPF conditions) with PAS638 *E. coli* (Figure 3A). PAS638 bacteria measured on at least 3 separate days over a >2-week period showed elevated tetrathionate sensing in IL10^−/−^ mice, but not separately-reared retired C57Bl/6 breeder controls (Figure 3B).

**Figure 3:**
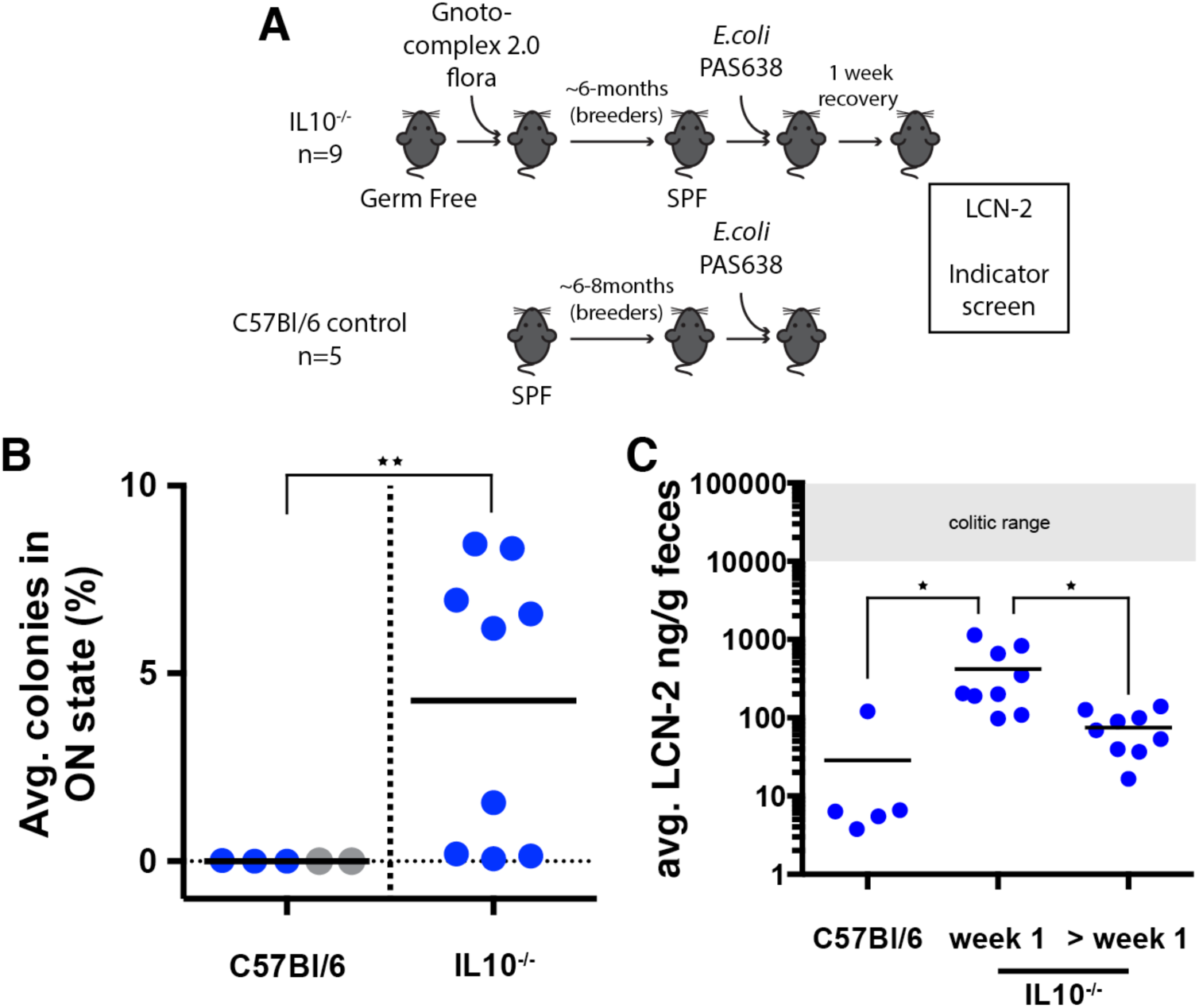
Tetrathionate levels are elevated in IL10^−/−^ mice. A) PAS638 was administered to retired breeder IL10^−/−^ mice raised in gnotobiotic and barrier SPF conditions and retired breeder C57Bl/6 control mice raised in SPF conditions. B) Indicator plating of fecal samples from C57Bl/6 and IL10^−/−^ mice (> 1 week post administration) showed elevated tetrathionate response in IL10^−/−^ mice. Blue values: average of 3-4 measurements from individual mice. Grey values: single measurement from individual mice. Means are marked ** p<0.01 between groups (including only blue values) using a Mann-Whitney test. C) LCN-2 levels were elevated in IL10^−/−^ compared to control mice in the week following PAS638 administration. Values are averages of 2-3 measurements from individual mice, with mean shown. *p< 0.05 using one-way ANOVA test with Tukey’s multiple comparison correction.

Switching levels in IL10^−/−^ mice were measured >1-week following bacterial administration to rule out inflammation caused by the process of administration, which included treatment with streptomycin - a known inflammatory enhancer in other colitis models ^25^. Indeed, fecal LCN-2 levels confirmed that inflammation in these mice was significantly elevated compared to controls in the week following bacterial administration but not subsequent to this when PAS638 switching was detected (Figure 3C).

Furthermore, these LCN-2 values were low compared to levels associated with colitic IL10^−/−^ mice ^22^. We conclude that ROS and tetrathionate are elevated in the IL10^−/−^ IBD model and that our engineered bacteria can sense this in a physiologically relevant subclinical inflammatory environment.

PAS638 maintains robust function over periods of at least 6-months colonizing the mammalian gut. An overarching issue in synthetic biology is that non-native genetic circuits may cause *in vivo* fitness costs resulting in mutation, loss-of-function and selective elimination from the host. To test genetic and functional stability we used the PAS638-inducing environment of 129X1/SvJ mice to maintain ON and OFF memory, thus testing fitness costs associated with both states. We administered PAS638 to mice and screened fecal samples for sensing at selected time-points over the following 200 days, > 1600 bacterial generations ^26^ (Figure 4A). PAS638 remained colonized at readily detectable titers throughout the experiment without further antibiotic selection (Figure 4B). As we observed previously (Figure 2B), tetrathionate levels as measured by PAS638 were variable in the 129X1/SvJ mice (Figure 4C). Consistent with the ability of PAS638 to detect inflammation, the presence of memory-ON state PAS638 bacteria broadly correlated with elevated LCN-2 levels, despite absolute LCN-2 levels being orders of magnitude lower than those measured during *Salmonella* infection (Figure S2D; Table S2). Furthermore, while it was not possible to quantify any overall inflammation by histology, an inflamed Peyer’s patch was observed from mouse 4 (data not shown).

**Figure 4:**
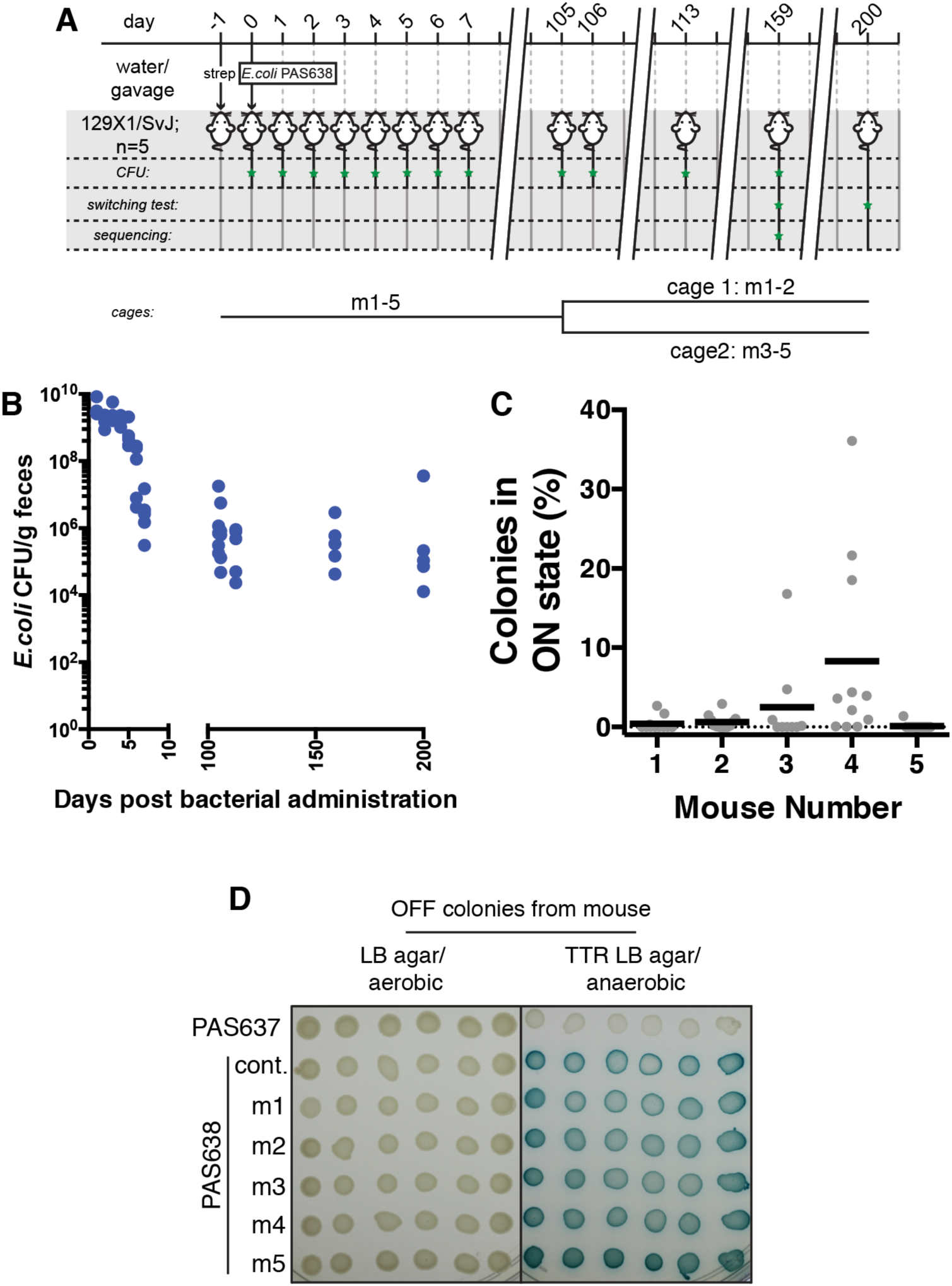
Tetrathionate-responsive memory circuits remain stable over >6 months colonizing the mammalian gut. A) 129X1/SvJ mice were colonized with PAS638 for a 200-day period. B) PAS638 titers remained steady following establishment in the streptomycin pre treated gut. C) Screening identified PAS638 response throughout the experiment. Graph shows switching of individual samples, with means. D) Colonies tested for function *in vitro* at day 159 showed tetrathionate specific switching.

Remarkably, colonies tested from fecal samples of mice at day 159 (Figure 4D) and day 200 (Table S3) retained the ability to respond to tetrathionate *in vitro*, indicating they had not acquired mutations to prevent their correct function.

Whole genome sequencing further demonstrated stability of these circuits. To analyze whether mutations had accrued in PAS638 during growth in the mammalian gut, mixtures of PAS638 colonies from each mouse at day 159 along with ancestral PAS638 were analyzed by whole genome sequencing. We sequenced to a depth capable of confidently identifying mutations greater than ~10% of the population (average chromosomal fold-coverage of 85-150). Only one unique mutation was identified from the five sequenced mixes (Table 1). This repeat expansion in an intergenic region of the *E. coli* NGF-1 colicin-like plasmid was detected in 89% and 92% of reads from two of the five mice (Table 1). Detection of a single mutant is consistent with the number expected from wild-type *E. coli* over >1200 generations ^27, 28^. The two mice carrying bacteria with the detected mutations were housed together, while all other mice were separated at day 105 (Figure 4A). Given this mutation spread and fixed amongst the cohoused mice, we are confident that any beneficial mutations would have been detected at the sequencing depth we achieved. Together these findings indicate that these synthetic circuits place sufficiently low burden on the bacteria carrying them to allow robust function over >6-months in the complex gut environment.

Here we used engineered bacteria to track a marker of the mammalian inflammatory response that was previously not amenable to non-invasive monitoring. Our results extend the previously limited knowledge of tetrathionate biology, confirming its association with inflammation in the presence and absence of pathogenic infection and specifically identifying its elevation in IL10^−/−^ mice and 129X1/SvJ mice (Figure 2B, 3B, Table S2). PAS638 proved sensitive to low-level inflammation in these models (Figure 3B-C; Table S2) and in some cases provided more consistent results over a multiple day period than LCN-2 levels (Table S2), possibly due to the memory capacity allowing signal integration over an extended period. The results point to a need for further research into this molecule, in particular whether it is similarly elevated in human inflammatory disease.

Several design features of this system stand out. The modularity of these circuits allows for inputs from any environmentally responsive promoter and outputs to any downstream gene circuit. Further, the self-regulating nature of the CI/Cro switch lowers the burden of these circuits on host cells. Together this establishes a platform that can be expanded for the non-invasive monitoring of other transient species in the gut, as well as the potential for easy future integration into more complex on-demand synthetic circuits, such as sense-and-respond therapeutics.

Areas for future investigation and improvement to this system remain. In particular, despite responding to low-levels of inflammation under certain settings, absolute PAS638 switching percentages were often variable and relatively low as a percentage of colonies screened. Several factors may contribute to this, including the sensitivity of the circuits, the time required in the presence of signal for switching to occur, the relative localization of bacteria compared inflammation and the excretion dynamics of the bacteria themselves.

Taken together, our work dispels the view that all synthetic gene circuits would be subject to mutational loss-of-function under extended growth in the complex, competitive setting of the mammalian gut and demonstrates the ability for synthetic bacterial devices to make unique impacts on our understanding of disease.

## Materials and Methods

### Strain construction

Details of strains constructed for this study are provided (Table S1). All synthetic constructs were integrated chromosomally in the mouse commensal *Escherichia coli* NGF-1 strain ^5^ or *Salmonella typhimurium* 14028s.

The lambda derived CI/Cro memory element was originally inserted between the *mhpR* and *lacZ* loci of *E. coli* TB10 ^29^ and transferred to *E. coli* NGF-1 by P1*vir* transduction ^30^ as previously described ^5^. The tetrathionate responsive trigger ^circuit, consisting of *ttrR, ttrS* genes and the P_ttrBCA_ promoter was amplified directly^ from *S. typhimurium* LT2. Overlap PCR was used to append the cro gene ^downstream of P_ttrBCA_ and add ~100bp flanks for insertion into the *araB-araC* locus^ of *E. coli* TB10. The locus was ultimately transferred to *E. coli* NGF-1 by P1*vir* transduction.

*S. typhimurium* strains were derived from *S. typhimurium* 14028s. To confer tetracycline resistance to all *S. typhimurium* strains a zhj-1401::Tn10 construct from *S. typhimurium* LT2 SA2700 (Salmonella Genetic Stock Center) transferred to both S. typhimurium strains used in this study by P22 transduction ^31^. To prevent tetrathionate reduction capacity, a *ttrR* knock-out TT22470 ^16^ construct was transferred to *S. typhimurium* by P22 transduction.

### *In vitro* induction with tetrathionate

Tetrathionate-based memory was induced under *in vitro* conditions in either liquid culture or during growth on plates. For liquid culture induction, strains were backdiluted 1:1000 from overnight culture in SOC broth into pre-reduced anaerobic SOC broth and grown at 37^o^C in an anaerobic chamber (Coy Lab Products) using 7%H_2_/ 20% CO_2_/63% N_2_ culture gas. After 1h, potassium tetrathionate (Sigma) dissolved in pre-reduced SOC media was added to cultures and induction was undertaken at 37^o^C in the anaerobic chamber for 4h. Memory was assayed by plating and growth in aerobic conditions of serial dilutions of the bacteria on luria-broth (LB) + 300μg/mL streptomycin sulfate (Sigma) + 60μg/mL 5-Bromo-4-chloro-3-indolyl b-D-galactopyranoside (X-gal) (Santa Cruz Biotechnology) agar plates.

Switching levels were estimated through counting greater than 250 blue and white colonies unless otherwise noted. *In vitro* testing on solid plates involved plating on LB + 300μg/mL streptomycin sulfate (Sigma) + 60μg/mL x-gal (Santa Cruz Biosciences) + 10mM sodium tetrathionate (Sigma) agar. Growth was maintained in anaerobic conditions for approximately 8-12 hours using the GasPak system (BD Biosciences), followed by further growth in aerobic conditions to allow development of memory and the x-gal reporter.

### *In vivo* testing of tetrathionate memory

The Harvard Medical School Animal Care and Use Committee approved all animal study protocols.

#### General analysis

For most experiments, female, Balb/c (Charles River), C57Bl/6 (Charles River) or 129X1/SvJ (Jackson laboratories) mice of 8-12 weeks, including ~2 weeks acclimatization to our mouse facility, were administered PAS638 *E. coli* NGF-1 bacteria (1-4 x 10^7^ CFU/ mouse) by oral gavage following pre-treatment with USP-grade streptomycin sulfate (Gold Biotechnology; 0.5g/L in drinking water or 20mg per mouse by oral gavage). Bacteria were pelleted and resuspended at a dilution of 1/10 from overnight culture in sterile PBS prior to gavage of 100uL/ mouse.

Fecal samples were collected from mice under study and homogenized at 50 or 100 mg/mL in sterile PBS by vortexing in 1.5mL eppendorf tubes for 5 minutes. Large debris was pelleted from homogenized feces by centrifugation at 200rpm (4g) for 20min in a benchtop centrifuge. Bacteria were cultured on agar plates following serial dilution of the resulting supernatant. Enumeration by colony counting and analysis of switching by comparison of blue (lac^+^, ON) and white (lac^−^, OFF) colony counts was achieved by plating on LB + 60μg/mL x-gal (Santa Cruz Biosciences) agar with 300μg/mL streptomycin sulfate (Sigma), 34 μg/mL chloramphenicol (Sigma) or a combination of both drugs. All mice were pre-screened for the presence of resistant colonies in feces pre bacterial administration. At least 250 colonies were counted per sample.

LCN-2 quantification was undertaken using the Mouse lipocalin-2/NGAL DuoSet ELISA kit (R&D Systems), using the manufacturer recommended protocol. Samples for ELISA were prepared as previously described ^22^, briefly, involving homogenization of 100mg/mL fecal pellets in PBS + 0.1% Tween20 by vortexing for 20 minutes at 4^°^C, followed by spinning at 12,000g for 10 minutes at 4^°^C. Clear supernatant was diluted at least 10 fold for quantification by ELISA. ELISA results were obtained on a Victor^3^V plate reader (Perkin Elmer) with 450/8nm and 540/8nm absorbance filters. For analysis, absorbance corrected values were interpolated from a sigmoidal four parameter logistic standard curve using Prism 6 for Mac OS X software (GraphPad).

At the conclusion of the experiments, where required, mice were sacrificed and dissected, their bowel removed and fixed whole in Bouin’s fixative (Sigma). Fixed tissues were embedded in paraffin, sectioned and stained with hematoxylin and eosin. Scoring of severity of inflammation was undertaken on single longitudinal sections of whole gut in a blinded fashion by a trained rodent histologist (RTB).

#### Streptomycin treated Salmonella colitis model

The streptomycin-treated Salmonella colitis model was undertaken in C57Bl/6 mice (Charles River) as previously described ^21^. Briefly, mice were administered with 20mg USP grade streptomycin sulfate (Gold Biotechnology) by oral gavage following 4h nil per os (NPO). 24 hours later, again following 4h NPO, mice were administered bacterial strains, resuspended from overnight culture in phosphate buffered saline (Gibco), by oral gavage. Bacteria were administered at ~1x10^7^ *E. coli* and ~1x10^8^ *S. typhimurium* per mouse as previously described ^21^

Selective plating for enumeration of *S. typhimurium* strains was achieved on M9 + 0.4% glucose + 30μg/mL tetracycline (Sigma) agar. Enumeration and analysis of switching was achieved using plating on LB + 300μg/mL streptomycin sulfate (Sigma) + 60μg/mL x-gal (Santa Cruz Biosciences) agar.

#### IL10 knockout model

Male retired breeder mice (~6 months age) from an IL10^−/−^ background ^32^ were transferred to barrier SPF conditions, following growth under gnotobiotic conditions. The mice were colonized with a complex set of human commensal microbes, Gnotocomplex 2.0 (Table S4), which is an expanded version of the Gnotocomplex designed to capture additional functionality and phylogenetic diversity ^33^. Control mice were male retired breeder mice (typically 6-8 months age) from a C57Bl/6 background raised in SPF conditions (Charles river). Mice were administered PAS638 bacteria (~3x10^7 per mouse) by oral gavage following USP streptomycin sulfate treatment (Gold Biotechnology; 0.5g/L in drinking water). In mice where colonization was lower than 2.5x10^4^ CFU/g following gavage, streptomycin was re-administered within the first week following bacterial administration for up to 48h to assist robust colonization.

Analysis of fecal samples was undertaken as described above. To avoid adverse influence of streptomycin and bacterial administration in IL10^−/-−^ mice, which are sensitive to aberrant inflammatory responses (as evident from LCN-2 quantification in week 1 post administration (Figure 4B)), blue-white colony screening was not undertaken in the week following administration. Results were generated from 3-4 fecal samples taken over the subsequent 2 weeks for control C57Bl/6 mice and ~1 month for IL10^−/−^ mice.

### Whole Genome Sequencing and analysis

Bacterial samples were selected by plating from feces, as described above, on LB + 300μg/mL streptomycin sulfate (Sigma) + 60μg/mL x-gal (Santa Cruz Biosciences) agar. Colonies were scraped from plates, resuspended and prepared for sequencing as a pool using a small-volume modification of the Illumina Nextera XP kit as described previously ^34^. Sequencing was performed on the Illumina Miseq platform, using paired end 50-bp reads.

Alignment and base calling of the ancestral sample was done via BreSeq ^35^ using default options, aligning to *E.coli* NGF-1 [Karrenbelt, 2016, in preparation; NCBI references NZ_CP016007.01, NZ_CP016008.01 and NZ_CP016009.01]. Breseq’s gdtools package was used to modify the ancestral genbank file to eliminate mutations in the ancestral strain. We then ran breseq at high sensitivity (settings-consensus-minimum-coverage-each-strand 3--consensus-frequency-cutoff 0.1) to call mutations in the passaged strains.

### Statistical analyses

All statistical analyses were undertaken in Prism 6 for Mac OS X (GraphPad). Details of individual tests and resulting p-values are included in figure legends and text where appropriate.

## Acknowledgements

We thank Amanda Graveline and Lynn Bry for discussions and assistance with mouse experiments. *S. typhimurium* TT22470 was a gift from John Roth. DTR is supported by a Human Frontier Science Program Long-term fellowship and an NHMRC/RG Menzies Postdoctoral fellowship. The research was funded by Defence Advanced Research Projects Agency Grant HR0011-15-C-0094 and the Wyss Institute for Biologically Inspired Engineering.

## References

1. Gardner, T.S., Cantor, C.R. & Collins, J.J. Construction of a genetic toggle switch in Escherichia coli. Nature 403, 339–342 (2000).

2. Archer, E.J., Robinson, A.B. & Süel, G.M. Engineered E. coli that detect and respond to gut inflammation through nitric oxide sensing. ACS Synth. Biol. 1, 451–457 (2012).

3. Yang, L. et al. Permanent genetic memory with >1-byte capacity. Nat. Methods 11, 1261–1266 (2014).

4. Farzadfard, F. & Lu, T.K. Synthetic biology. Genomically encoded analog memory with precise in vivo DNA writing in living cell populations. Science 346, 1256272 (2014).

5. Kotula, J.W. et al. Programmable bacteria detect and record an environmental signal in the mammalian gut. Proceedings of the National Academy of Sciences of the United States of America 111, 4838–4843 (2014).

6. Mimee, M., Tucker, A.C., Voigt, C.A. & Lu, T.K. Programming a Human Commensal Bacterium, Bacteroides thetaiotaomicron, to Sense and Respond to Stimuli in the Murine Gut Microbiota. Cell Syst 1, 62–71 (2015).

7. Mosli, M.H. et al. C-Reactive Protein, Fecal Calprotectin, and Stool Lactoferrin for Detection of Endoscopic Activity in Symptomatic Inflammatory Bowel Disease Patients: A Systematic Review and Meta-Analysis. Am J Gastroenterology 110, 802–819-quiz 820 (2015).

8. Bouguen, G. et al. Treat to target: a proposed new paradigm for the management of Crohn’s disease. Clin. Gastroenterol. Hepatol. 13, 1042–1050.e1042 (2015).

9. Sipponen, T., Nuutinen, H., Turunen, U. & Färkkilä, M. Endoscopic evaluation of Crohn's disease activity: comparison of the CDEIS and the SES-CD. Inflamm. Bowel Dis. 16, 2131–2136 (2010).

10. Travis, S.P.L. et al. Reliability and initial validation of the ulcerative colitis endoscopic index of severity. Gastroenterology 145, 987–995 (2013).

11. Brazil, J.C., Louis, N.A. & Parkos, C.A. The role of polymorphonuclear leukocyte trafficking in the perpetuation of inflammation during inflammatory bowel disease. Inflamm. Bowel Dis. 19, 1556–1565 (2013).

12. Winter, S.E. et al. Gut inflammation provides a respiratory electron acceptor for Salmonella. Nature 467, 426–429 (2010).

13. Kamdar, K. et al. Genetic and Metabolic Signals during Acute Enteric Bacterial Infection Alter the Microbiota and Drive Progression to Chronic Inflammatory Disease. Cell Host Microbe 19, 21–31 (2016).

14. Hensel, M., Hinsley, A.P., Nikolaus, T., Sawers, G. & Berks, B.C. The genetic basis of tetrathionate respiration in Salmonella typhimurium. Molecular microbiology 32, 275–287 (1999).

15. Winter, S.E., Lopez, C.A. & Bäumler, A.J. The dynamics of gut-associated microbial communities during inflammation. EMBO Rep 14, 319–327 (2013).

16. Price-Carter, M., Tingey, J., Bobik, T.A. & Roth, J.R. The alternative electron acceptor tetrathionate supports B12-dependent anaerobic growth of Salmonella enterica serovar typhimurium on ethanolamine or 1,2-propanediol. J. Bacteriol. 183, 2463–2475 (2001).

17. Liu, Y.-W., Denkmann, K., Kosciow, K., Dahl, C. & Kelly, D.J. Tetrathionate stimulated growth of Campylobacter jejuni identifies a new type of bi-functional tetrathionate reductase (TsdA) that is widely distributed in bacteria. Molecular microbiology 88, 173–188 (2013).

18. Rooks, M.G. et al. Gut microbiome composition and function in experimental colitis during active disease and treatment-induced remission. ISME J 8, 1403–1417 (2014).

19. White, P., Liebhaber, S.A. & Cooke, N.E. 129X1/SvJ mouse strain has a novel defect in inflammatory cell recruitment. The Journal of Immunology 168, 869–874 (2002).

20. Hoover-Plow, J.L., et al. Strain and model dependent differences in inflammatory cell recruitment in mice. Inflamm. Res. 57, 457–463 (2008).

21. Barthel, M. et al. Pretreatment of mice with streptomycin provides a Salmonella enterica serovar Typhimurium colitis model that allows analysis of both pathogen and host. Infect. Immun. 71, 2839–2858 (2003).

22. Chassaing, B. et al. Fecal lipocalin 2, a sensitive and broadly dynamic non-invasive biomarker for intestinal inflammation. PLoS ONE 7, e44328 (2012).

23. Simmonds, N.J. & Rampton, D.S. Inflammatory bowel disease--a radical view. Gut 34, 865–868 (1993).

24. Keubler, L.M., Buettner, M., Häger, C. & Bleich, A. A Multihit Model: Colitis Lessons from the Interleukin-10-deficient Mouse. Inflamm. Bowel Dis. 21, 1967–1975 (2015).

25. Spees, A.M. et al. Streptomycin-induced inflammation enhances Escherichia coli gut colonization through nitrate respiration. MBio 4 (2013).

26. Myhrvold, C., Kotula, J.W., Hicks, W.M., Conway, N.J. & Silver, P.A. A distributed cell division counter reveals growth dynamics in the gut microbiota. Nat Commun 6, 10039 (2015).

27. Drake, J.W., Charlesworth, B., Charlesworth, D. & Crow, J.F. Rates of spontaneous mutation. Genetics 148, 1667–1686 (1998).

28. Lee, H., Popodi, E., Tang, H. & Foster, P.L. Rate and molecular spectrum of spontaneous mutations in the bacterium Escherichia coli as determined by whole-genome sequencing. Proceedings of the National Academy of Sciences of the United States of America 109, E2774–2783 (2012).

29. Datsenko, K.A. & Wanner, B.L. One-step inactivation of chromosomal genes in Escherichia coli K-12 using PCR products. Proceedings of the National Academy of Sciences of the United States of America 97, 6640–6645 (2000).

30. Miller, J.H. Experiments in molecular genetics. (Cold Spring Harbor Laboratory, 1972).

31. Davis, R.W., Botstein, D. & Roth, J.R. A Manual for Genetic Engineering: Advanced Bacterial Genetics. (Cold Spring Harbor Laboratory, 1980).

32. Devkota, S. et al. Dietary-fat-induced taurocholic acid promotes pathobiont expansion and colitis in Il10-/- mice. Nature 487, 104–108 (2012).

33. Bucci, V. et al. MDSINE: Microbial Dynamical Systems INference Engine for microbiome time-series analyses. Genome Biol. 17, 121 (2016).

34. Baym, M. et al. Inexpensive multiplexed library preparation for megabase-sized genomes. PLoS ONE 10, e0128036 (2015).

35. Deatherage, D.E. & Barrick, J.E. Identification of mutations in laboratory-evolved microbes from next-generation sequencing data using breseq. Methods Mol. Biol. 1151, 165–188 (2014).

